# Emergence of resistance to the antiparasitic selamectin in *Mycobacterium smegmatis* is improbable and contingent on cell wall integrity

**DOI:** 10.1101/2024.09.05.611451

**Authors:** José Manuel Ezquerra-Aznárez, Henrich Gašparovič, Álvaro Chiner-Oms, Ainhoa Lucía, Jesús Blázquez, Iñaki Comas, Jana Korduláková, José A. Ainsa, Santiago Ramón-García

## Abstract

Tuberculosis remains the deadliest infectious disease of the 21^st^ century. New antimicrobials are needed to improve treatment outcomes and enable therapy shortening. Drug repurposing is an alternative to the traditional drug discovery process. The avermectins are a family of macrocyclic lactones with anthelmintic activity active against *Mycobacterium tuberculosis*. However, their mode of action in mycobacteria remains unknown.

In this study, we employed traditional mutant isolation approaches using *Mycobacterium smegmatis*, a non-pathogenic *Mycobacterium tuberculosis* surrogate. We were only able to isolate mutants with decreased susceptibility to selamectin using the Δ*nucS* mutator *M. smegmatis* strain. This phenotype was caused by mutations in *mps1* and *mmpL11*. Two of these mutants were used for a second experiment in which high-level selamectin-resistant mutants were isolated; however, specific mutations driving the phenotypic change to high-level resistance could not be identified. The susceptibility to selamectin in these mutants was restored to the basal level by subinhibitory concentrations of ethambutol. The selection of ethambutol resistance in a high-level selamectin-resistant mutant also resulted in multiple colonies becoming susceptible to selamectin again. These colonies carried mutations in *embB*, suggesting that integrity of the cell envelope is a prerequisite for selamectin resistance. The absence of increased susceptibility to selamectin in an *embB* deletion strain demonstrated that the target of selamectin is not cytosolic. Our data show that concurrence of specific multiple mutations and complete integrity of the mycobacterial envelope are necessary for selamectin resistance. Our studies provide first-time insights into the antimycobacterial mode of action of the antiparasitic avermectins.

## Introduction

Tuberculosis (TB) is the leading cause of death among infectious diseases during the 21^st^ century. In 2022 an estimated 1.3 million people died from TB, and 10.6 million developed the disease. Despite being a curable disease, the global TB situation has been aggravated in the last decades by the emergence of drug-resistant strains, which currently account for about 3% of the newly diagnosed cases and about 20% of the relapses [1]. The recent introduction of novel and repurposed anti-TB drugs has allowed the development of shorter and more effective treatments for drug-resistant strains [2–4]. However, drug-resistant TB remains the first cause of death due to antimicrobial resistance, and the overall treatment success for drug-resistant strains remains below 60% [5,6]. New anti-TB drugs are thus needed to improve treatment success and prevent the emergence of resistance to the recently introduced compounds.

Developing a new antimicrobial from the bench to the clinic requires enormous time and economic investments, which could take more than a decade and up to more than 2 billion USD [7]. Moreover, antimicrobials are often seen as drugs with reduced profitability, discouraging pharmaceutical companies from investing in antimicrobial discovery [8]. In recent years, drug repurposing, i.e., finding new applications for clinically approved drugs, is gaining momentum as an alternative to the *de novo* discovery process [9]. It is estimated that around 4,000 approved drugs are being repurposed for new medical applications [10]. Using this approach, it was found that avermectins are active against *Mycobacterium tuberculosis*, the causing agent of TB, and other pathogenic mycobacteria [11–13]. This was an unexpected finding given their lack of activity against Gram-positive and Gram-negative bacteria [11].

Avermectins are a family of 16-membered macrocyclic lactones with broad-spectrum anthelmintic activity. They were isolated from *Streptomyces avermitilis* in the 1970s. Ivermectin, a semi-synthetic derivative of the original avermectins, was approved for use in humans in 1987 [14,15] and it has been used ever since in the prevention of onchocerciasis and lymphatic filariasis [16]. Recent studies have identified potential new activities: ivermectin can control glucose and cholesterol levels in diabetic mice, suppress the proliferation of cells in various cancer models, and inhibit the replication of several viruses. Furthermore, ivermectin has been proposed as a tool to prevent the transmission of vector-borne diseases, such as malaria [17,18]. Other avermectins are widely used to treat parasite infections in livestock and pets. The antiparasitic mechanism of action of avermectins is well-known: avermectins bind to glutamate-gated chloride channels in muscle and nerve cells and increase the permeability to chloride ions. This causes membrane hyperpolarization and eventually leads to death by paralysis [19]. Their mode of action against mycobacteria, however, remains unknown.

In a previous study, we demonstrated the *in vitro* inhibitory activity of selamectin against the decaprenylphosphoryl-β-D-ribose oxidase (DprE1). However, subsequent studies using whole cells showed that DprE1 is not the sole target of selamectin in mycobacteria, suggesting that avermectins might interact with multiple targets [20].

In this study, we isolated and characterized selamectin-resistant mutants in *Mycobacterium smegmatis*, a non-pathogenic fast-growing surrogate of *M. tuberculosis*, and identified essential requirements needed for intermediate and high-level resistance to selamectin in mycobacteria.

## Results

### Characterization of selamectin activity against *M. smegmatis*

To establish optimal conditions for mutant isolation and subsequent characterization of selamectin-resistant colonies, we determined the activity of selamectin against the model strain *M. smegmatis* mc^2^155 in liquid and solid media. The MIC of selamectin was 4 µg/mL in all media tested, except the one containing the detergent tyloxapol. Because of this, the use of detergents was discarded for assays involving selamectin. Selamectin activity was tested next against different inoculums of *M. smegmatis* in solid 7H10-OADC agar medium. The MIC of selamectin showed a strong inoculum dependence (**Table 1**).

**Table 1.** MIC of selamectin against *M. smegmatis* in different media.

| Broth Medium | MIC (µg/mL) |
| --- | --- |
| 7H9-0.2% glycerol-ADC | 4 |
| 7H9-0.2% glycerol-ADC-0.05% tyloxapol | 64 |
| LB | 4 |
| NE | 4 |
| Müller-Hinton II | 4 |
| Agar Medium<sup>a</sup> | MIC (µg/mL) |
| 10<sup>3</sup> | 4 |
| 10<sup>5</sup> | 8 |
| 10<sup>7</sup> | 16-32 |

<sup>a</sup>MIC in solid medium was tested on 7H10-OADC agar. The activity of selamectin was tested against different starting inoculums (10<sup>3</sup>, 10<sup>4</sup>, 10<sup>5</sup> CFU)

### Selamectin resistance cannot be selected by traditional mutant isolation approaches in a wild-type genetic background

We first unsuccessfully attempted traditional mutant isolation assays using the wild-type *M. smegmatis* mc^2^155 strain, with two different outcomes: inoculums below 10^7^ CFU yielded no colonies at selamectin concentrations higher than 4x MIC, whereas higher inoculums resulted in the growth of bacterial lawns. None of the colonies isolated from the borders of those plates displayed changes in selamectin susceptibility after being propagated in selamectin-free medium, suggesting that their isolation was due to a transient phenotype. We observed comparable outcomes when the inoculum was a pool of mutants of a transposon library made in *M. smegmatis* HS42 using the TnSPAZ transposon [21] and similar experimental setups. Fifty colonies were tested for susceptibility changes, and none displayed an increased MIC to selamectin. This suggested that no stable genetic mutations associated with selamectin resistance could be selected in a wild-type background. However, internal control mutant isolation assays carried in parallel using 4x MIC isoniazid allowed for the selection of stable resistant mutants with a mutant frequency of 1.8·10^-5^, which was comparable to previous reports [22].

### Low-level selamectin-resistant mutants can be selected in a mutator genetic background

We subsequently tried to isolate selamectin-resistant mutants using the *M. smegmatis* Δ*nucS* strain, which lacks a functional DNA mismatch repair system and shows mutant frequencies ca. 100-fold higher than wild-type strains [23]. We were able to isolate 38 colonies from the mutator strain, while no colonies were recovered in a parallel assay run with the reference *M. smegmatis* mc^2^155. The mutant frequency was 1.0·10^-6^, whereas isoniazid displayed a mutant frequency of 5.2·10^-4^ at 4x MIC (**Table S1**).

We subsequently determined the MIC of the 38 colonies and found that two of them displayed a 2-fold increase and 23 had a 1.4-fold increase in their MIC to selamectin. Despite being modest variations, results were consistent across several independent assays. Thus, we performed additional validation studies using drop-dilution (**Fig. 1A**), and time-kill kinetics (**Fig. 1B**) assays. Both assays correlated well: clones displaying increased survival on selamectin-containing agar plates were also less susceptible to selamectin on time-kill kinetic assays, thus confirming the subtle susceptibility changes observed.

**Fig. 1.**
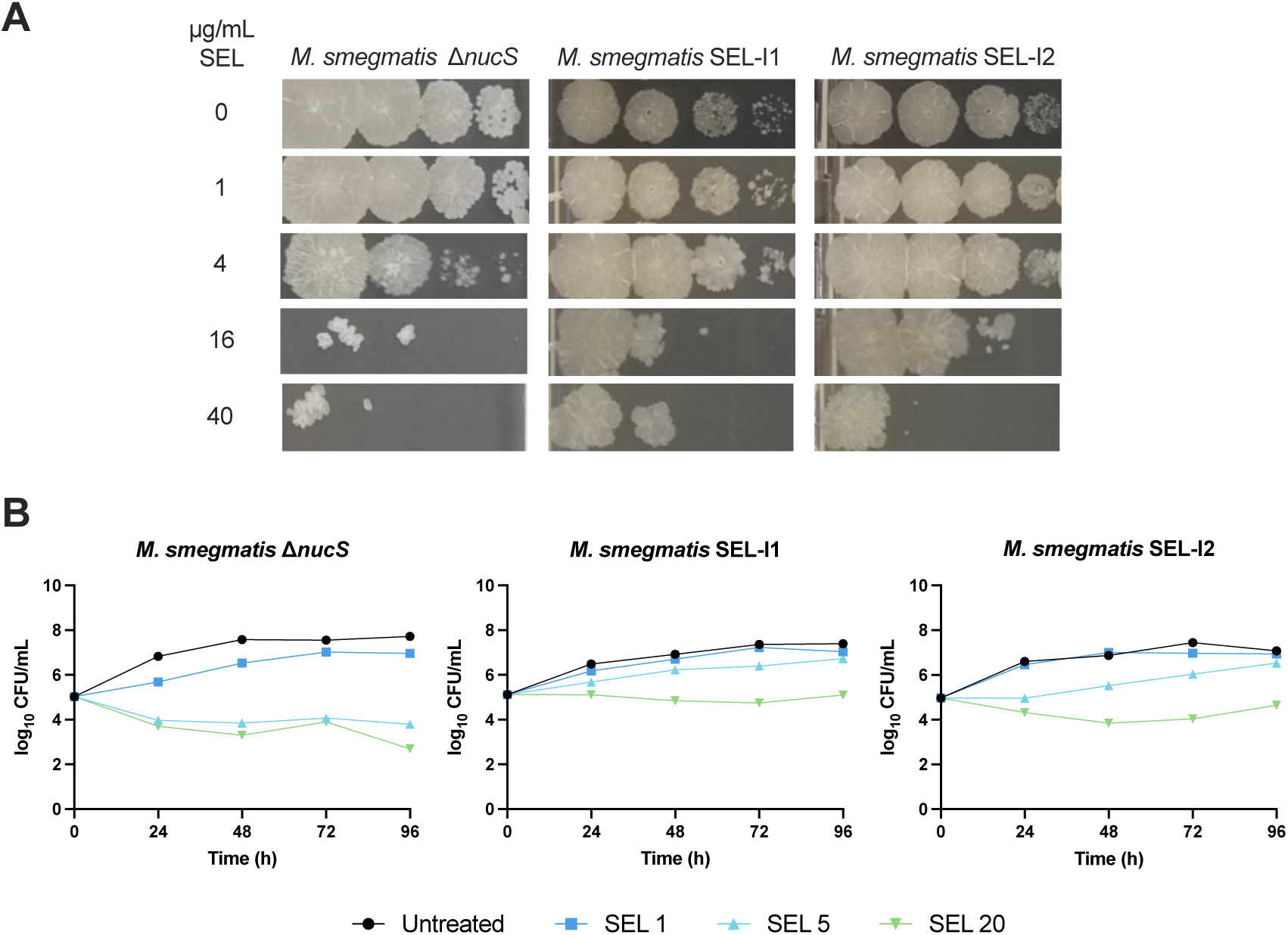
Selamectin resistance in mutants isolated from the mutator strain *M. smegmatis* Δ*nucS*. Both SEL-I1 and SEL-I2 carry nonsense mutations in *mps1*, SEL-I1 carries an additional nonsense mutation in *mmpL11*. (A). Drop dilution assay 10-fold serial dilutions of the bacterial inoculum were made from left to right. (B). Time-kill kinetics assay. Selamectin (SEL) concentrations are expressed in µg/mL.

### Concurrence of nonsense mutations in *mps1* and *mmpL11* is necessary for intermediate selamectin resistance

We then performed whole genome sequencing (WGS) of 10 selected clones (referred to as SEL-I1 to SEL-I10), which revealed the presence of a nonsense mutation in *mps1* (*MSMEI_0391*) in 5 out of the 10 clones. One of them harbored an additional nonsense mutation in the *mmpL11* (*MSMEI_0234*) locus. Additional non-synonymous mutations are shown in **Table S2**.

Given the sheer number of mutations identified, genetic validation efforts were focused on the two nonsense mutations. Single-stranded DNA recombineering was used to introduce them in the parental *M. smegmatis* Δ*nucS* strain. The reason to use this strain was the bias of NucS toward the correction of G-T, T-T, and G-G mismatches [24], which were predominant among the SNPs found in the sequenced mutants. The first recombineering attempt to reproduce the two nonsense mutations was carried out using selamectin as the selectable marker. However, no colonies with the desired mutations were retrieved when using ssDNA substrates encoding the nonsense mutations. This observation aligned with the results from mutant isolation assays carried out with the wild-type *M. smegmatis* mc^2^155, in which plating large inoculums resulted in the isolation of colonies without stable resistance phenotypes to selamectin.

Thus, selamectin selection was replaced by streptomycin (Sm) resistance as co-selection. This approach allowed the introduction of the *mps1* and *mmpL11* nonsense mutations in the *M. smegmatis* Δ*nucS* background. Whereas only the *mmpL11* engineered strain displayed a subtle increase in the MIC of selamectin in broth, both mutants displayed slight decreased susceptibility to selamectin in drop dilution assays (**Fig. 2** and **Table 2**).

**Fig. 2.**
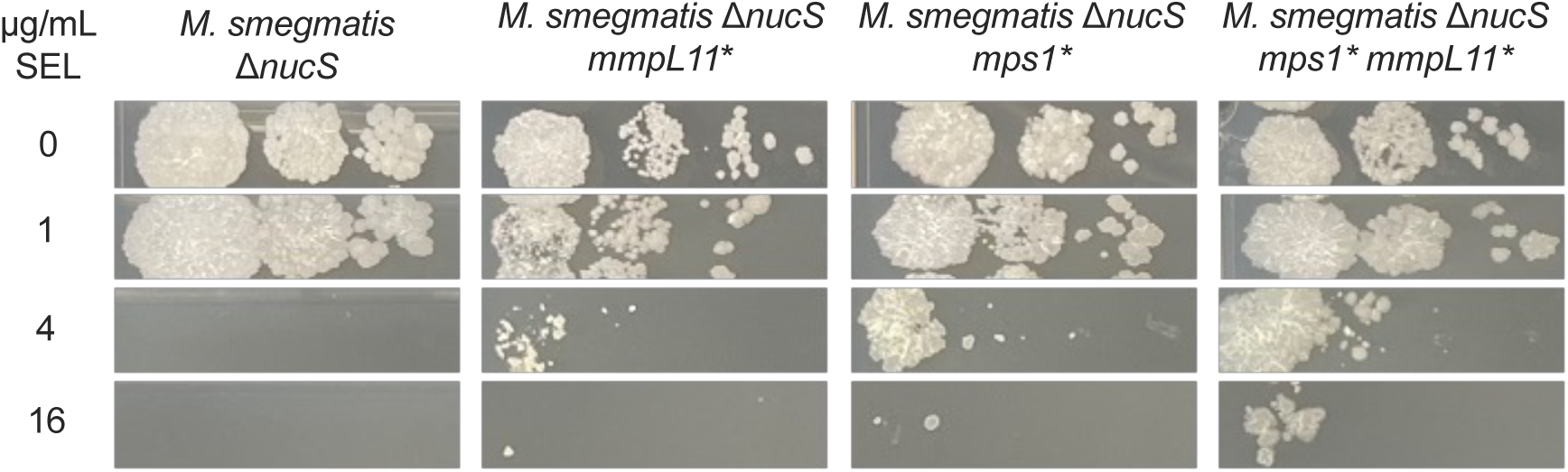
Phenotypic validation of *M. smegmatis* engineered point mutants at *mps1* and *mmpL11* by drop dilution assays. Selamectin (SEL) concentrations are expressed in µg/mL. 10-fold serial dilutions of the bacterial inoculum were made from left to right. *: truncated protein, due to a stop codon in the ORF.

**Table 2.** MIC of selamectin for *M. smegmatis* Δ*nucS* engineered strains.

| Strain | MIC (μg/mL) |
| --- | --- |
| *M. smegmatis* ΔnucS | 4 |
| *M. smegmatis* ΔnucS mps1* | 4 |
| *M. smegmatis* ΔnucS mmpL11* | 5.6 |
| *M. smegmatis* ΔnucS mps1*, mmpL11* | 5.6 |
| *M. smegmatis* ΔnucS mshA* | 5.6 |
| *M. smegmatis* ΔnucS MSMEI_1249* | 5.6 |
| *M. smegmatis* ΔnucS mps1*, mmpL11*, MSMEI_1249*, mshA* | 5.6 |

A double mutant carrying both nonsense mutations was built in order to assess whether these mutations had an additive effect on selamectin resistance. The coexistence of both mutations reproduced the phenotype observed in the intermediate SEL-I1 mutant, and the double mutant was less susceptible compared to each of the single mutants (**Fig. 2**).

### High-level resistant mutants to selamectin require mutations in different loci

Following the genetic validation of *mps1* and *mmpL11* mutations, a new mutant isolation assay was set using two of the previously isolated mutants: SEL-I1 (carrying nonsense mutations in *mps1* and *mmpL11*, among other point mutations) and SEL-I2 (carrying the nonsense mutation in *mps1*, among other point mutations). The MIC of selamectin was tested in 10 colonies derived from each mutant, displaying MIC > 32 µg/mL. This MIC shift was confirmed by time-kill kinetics in 9 out of the 10 total mutants tested (**Fig. 3** and **Fig. S1**).

**Fig. 3.**
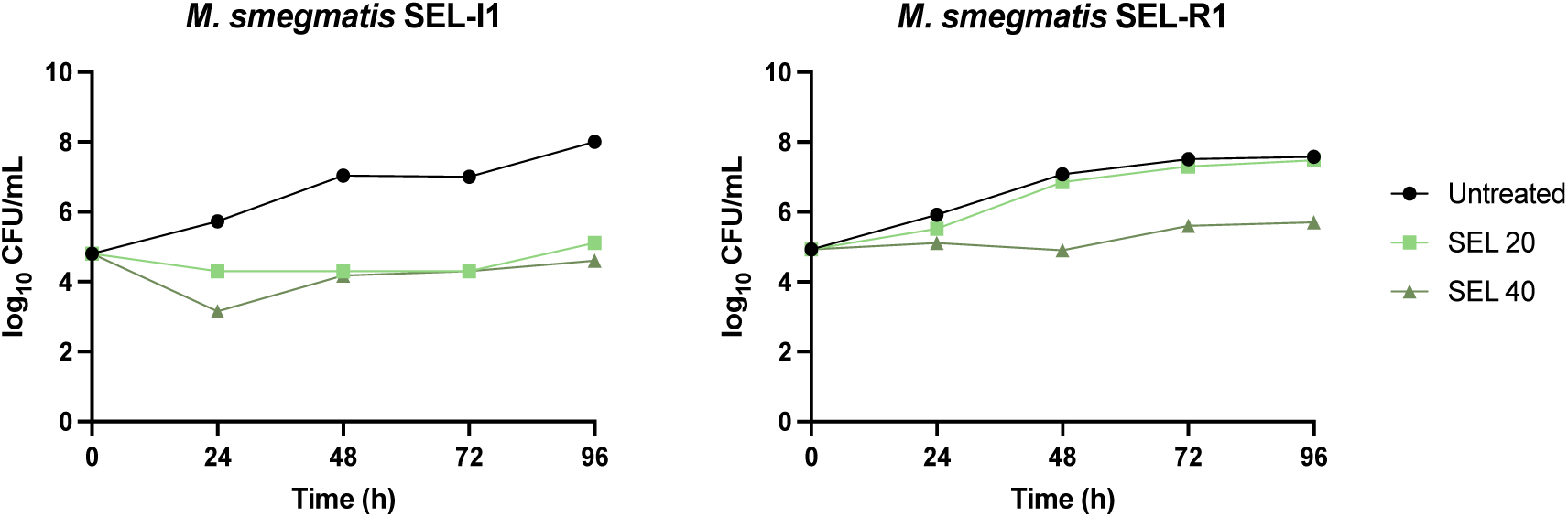
Time-kill kinetics of selamectin against SEL-R1, a representative high-level selamectin-resistant clone, and its parental strain (SEL-I1). Selamectin (SEL) concentrations are given in µg/mL.

WGS identified differences between the two genetic backgrounds. On the one hand, the four clones isolated from SEL-I1 (named SEL-R1 to SEL-R4) carried identical mutations, suggesting they derived from a single clone (**Table S3**), with two additional nonsense mutations in *mshA* and *MSMEI_1249*. On the other hand, the five clones isolated from SEL-I2 (named SEL-R5 to SEL-R9) carried different mutations from each other (**Table S4** and **Table S5**). Four of them shared a missense mutation in *mmpL11*, which confirmed the role of this locus in the acquisition of selamectin resistance and that *mps1* mutations were likely the first step in the process of acquiring resistance. Another mutation shared by several clones isolated from SEL-I2 was the substitution of the *mspA* canonical start codon to the less efficient GUG.

Mutations in *mshA* and *MSMEI_1249* were introduced separately in the *M. smegmatis* Δ*nucS* background. Neither of them reproduced the high-level selamectin resistance (**Fig. S2**). Then, both mutations were introduced sequentially in the engineered strain carrying *mps1* and *mmpL11* nonsense mutations. Similarly, they had no effect on the susceptibility to selamectin, as the increased survival observed at 4 µg/mL was due to the previously introduced mutations responsible for the SEL-I phenotype (**Fig. S2** and **Table 2**).

In order to study the role of porins in the susceptibility of *M. smegmatis* to selamectin (identified in SEL-I2, *mspA*), the MIC of selamectin was determined against porin-deficient *M. smegmatis* strains. We speculated that the substitution identified at the start codon would reduce the amount of protein; however, porin-deficient mutants remained susceptible to selamectin as the parental strain (**Table S6**).

### Selamectin-resistant mutants display altered composition of glycopeptidolipids

WGS of selected intermediate resistant strains (SEL-I1 and SEL-I2) and derived high-level resistant mutants (R1-R9) revealed the presence of nonsense mutations in genes *mps1* and *mmpL11*, both involved in the biogenesis and integrity maintenance of mycobacterial cell envelope. In *M. smegmatis*, *mps1* encodes a non-ribosomal peptide synthase playing a role in the biosynthesis of glycopeptidolipids (GPLs), and *mmpL11* is described as a part of lipid export machinery [25–29]. To confirm whether the aforementioned mutations induce alterations in the cell envelope of tested *M. smegmatis* strains, total lipids, alkaline-stable GPLs, and fatty/mycolic acids were extracted and analyzed from wild-type and mutant strains grown in 7H9 broth at 37 °C. Compared to wild-type, TLC analyses did not reveal significant changes in composition of mycolic and fatty acids, mycolic acids-derived molecules, phosphatidylethanolamine, cardiolipin, triacylglycerols or phosphatidylinositol mannosides in selamectin-resistant strains. However, in correlation with results obtained by WGS, resistant strains carrying mutations in gene *mps1* or in genes *mps1*/*mmpL11* displayed notable alterations in the production of GPLs (**Fig. 4E**).

**Fig. 4.**
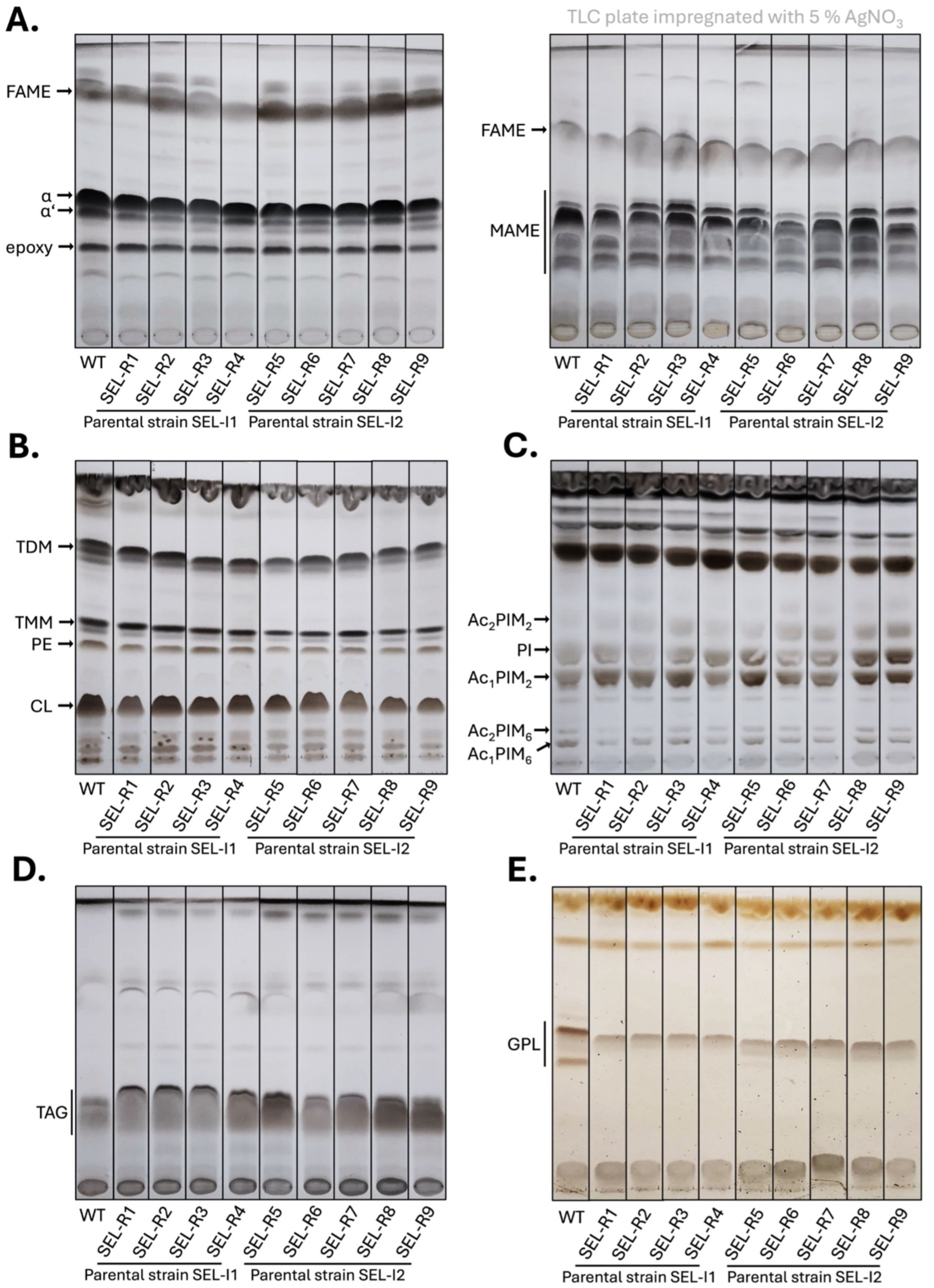
Thin-layer chromatography analysis of mycolic/fatty acids and lipids isolated from selamectin-resistant strains of *M. smegmatis*. (A) Thin-layer chromatography (TLC) analyses of mycolic and fatty acids methyl esters isolated from whole cells. Samples were loaded on standard (left) or AgNO_3_ impregnated (right) TLC plates and separated in solvent V. Isolated lipids were separated in **(B)** solvent I; **(C)** solvent II and **(D)** solvent III. **(E)** Glycopeptidolipids were separated in solvent IV. Lipids and mycolic/fatty acids were visualized with 10 % CuSO_4_ in 8 % phosphoric acid solution and charring. Glycopeptidolipids were visualized with orcinol in H_2_SO_4_ followed by charring. FAME, fatty acids methyl esters; MAME, mycolic acids methyl esters; *ɑ*, *ɑ*’ and epoxy refer to forms of mycolic acids methyl esters; TDM, trehalose dimycolates; TMM, trehalose monomycolates; PE, phosphatidylethanolamine; CL, cardiolipin; Ac_2_PIM_2_, diacylated phosphatidylinositol dimannosides; PI, phosphatidylinositol; Ac_1_PIM_2_, monoacylated phosphatidylinositol dimannosides; Ac_2_PIM_6_, diacylated phosphatidylinositol hexamannosides; Ac_1_PIM_6_, monoacylated phosphatidylinositol hexamannosides; TAG, triacylglycerols; GPL, glycopeptidolipids.

### Subinhibitory concentrations of cell wall inhibitors restored susceptibility to selamectin in the SEL-R1 background

Intermediate selamectin resistance appears to be driven by mutations that affect cell wall composition, and those changes seem necessary to develop high-level resistance. We then hypothesized that the combination of selamectin with cell wall targeting compounds could restore the susceptibility to this drug in the mutant background. To validate this hypothesis, the MIC of selamectin against the mutant SEL-R1 and the wild-type strain *M. smegmatis* mc^2^155 was determined in the presence of subinhibitory concentrations (0.25x MIC) of compounds targeting peptidoglycan (vancomycin, D-cycloserine, imipenem, cefradine, amoxicillin), arabinogalactan (ethambutol, BTZ043), and mycolic acid (isoniazid) biosynthesis. Compounds could be divided into three groups based on their effect on the MIC of selamectin in both strains (**Table 3**). First, most compounds showed no interaction with selamectin (i.e., the MIC of selamectin did not change in their presence of subinhibitory concentrations of these compounds). The second group (rifampicin, kanamycin) produced similar fold changes in the MIC in both strains, suggesting that their interaction with selamectin was independent of the mutations in the SEL-R1 strain. A third group, represented by ethambutol, vancomycin, and cefradine, showed a differential interaction with the SEL-R1 background, which resulted in restoration of the susceptibility to selamectin.

**Table 3.** Variations in the MIC of selamectin in the presence of 0.25x MIC of cell wall inhibitors. *Fold change is calculated as the ratio between the MIC of SEL alone divided by the MIC of SEL in the presence of 0.25x MIC of a given compound. The MIC of selamectin was 2 µg/mL for *M. smegmatis* mc^2^155, and 8-16 µg/mL for *M. smegmatis* SEL-R1. Kanamycin and rifampicin were included as controls because they target processes that are not directly related to the cell wall.

| Compound | Fold change*<br>(*M. smegmatis* mc<sup>2</sup>155) | Fold change*<br>(*M. smegmatis* SEL-R1) |
| --- | --- | --- |
| Cefradine | 2 | 16 |
| Ethambutol | 1 | 8 |
| Vancomycin | 4 | 8 |
| D-cycloserine | 1 | 1 |
| Imipenem | 1 | 1 |
| Isoniazid | 1 | 1 |
| Amoxicillin | 1 | 1 |
| BTZ043 | 1 | 1 |
| Rifampicin | 4 | 4 |
| Kanamycin | 2 | 2 |

We then studied in more detail the interaction between selamectin and the first-line anti-TB drug ethambutol. For the reference strain *M. smegmatis* mc^2^155, the MIC of selamectin did not change in the presence of ethambutol, and no interaction was detected (FICI = 0.75). In contrast, in the SEL-R1 mutant background the interaction between selamectin and ethambutol was confirmed with a FICI of 0.25, indicative of synergism (**Fig. 5**).

**Fig. 5.**
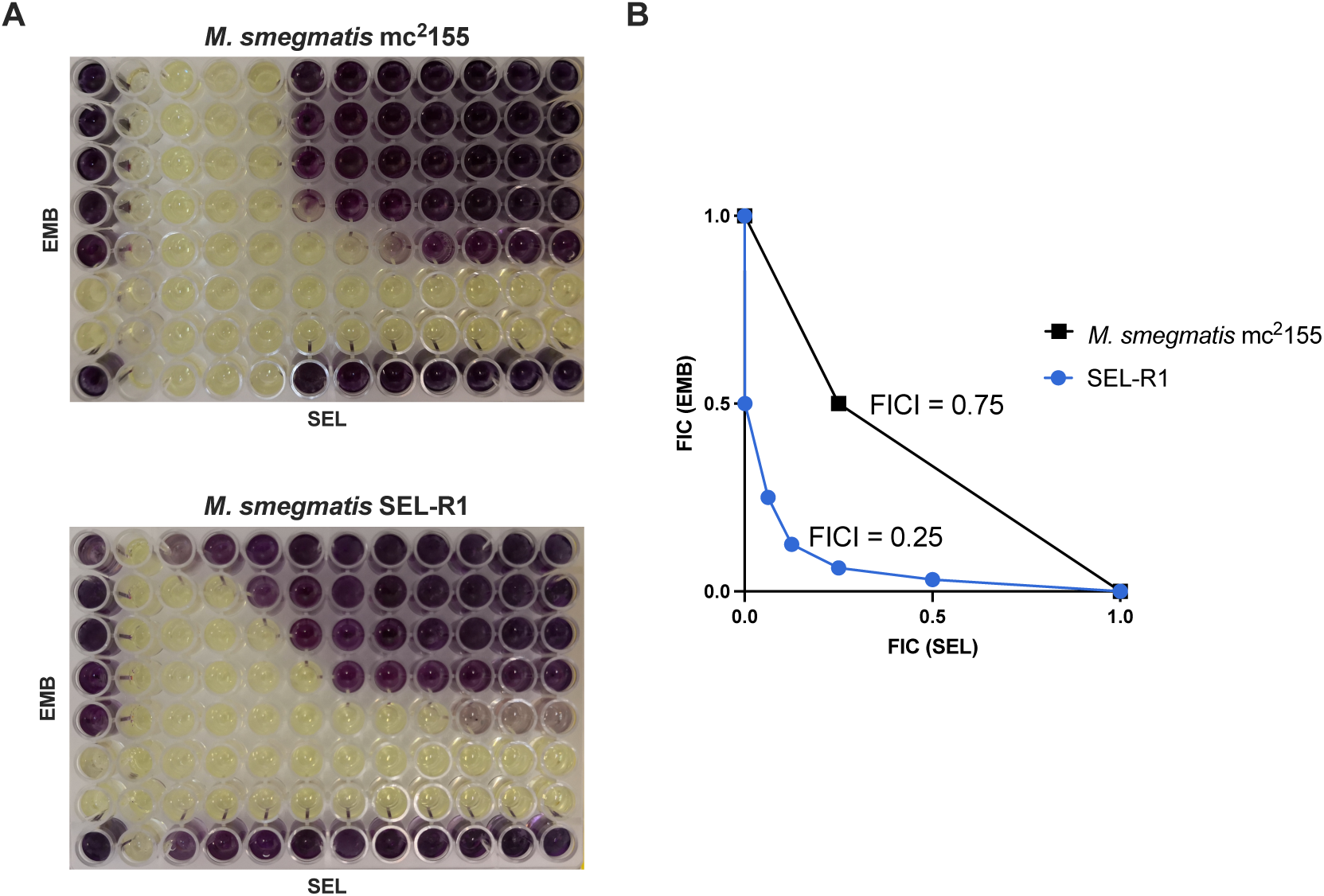
Selamectin and ethambutol interaction studies in *M. smegmatis*. (A) Wild-type *M. smegmatis* mc^2^155 and mutant *M. smegmatis* SEL-R1 strains were tested by the checkerboard assay. Dark violet wells indicate bacterial growth, while yellow wells, growth inhibition. Maximum concentrations used were 4 µg/mL for ethambutol and 32 µg/mL for selamectin. (B) Isobologram plot for the selamectin-ethambutol interaction in each genetic background.

Ethambutol was thus able to restore selamectin wild-type susceptibility levels in a selamectin-resistant mutant. We speculated whether the opposite effect (i.e., selamectin reducing the MIC of ethambutol in an ethambutol-resistant mutant) could also be possible. For this, ethambutol-resistant mutants were generated from two different backgrounds: (i) the *M. smegmatis* Δ*nucS* (selamectin-susceptible) and (ii) the SEL-R1 selamectin-resistant mutant (a derivative of *M. smegmatis* Δ*nucS*). Sanger sequencing revealed that all fourteen ethambutol-resistant mutants selected—6 from *M. smegmatis* Δ*nucS* and 8 from *M. smegmatis* SEL-R1—carried mutations in *embB*, although at different positions (**Table 4**). We then tested the activity of selamectin against the newly generated *embB* mutants. Susceptibility to selamectin in the mutants derived from *M. smegmatis* Δ*nucS* did not change compared to the parental strain. However, acquisition of ethambutol resistance in the SEL-R1 background resulted in decreased MICs of selamectin in 6 of the 8 colonies tested (**Table 4**). Time-kill kinetics of four strains representing all possible combinations of susceptibility to selamectin and ethambutol showed that subinhibitory concentrations of ethambutol could make selamectin subMIC concentrations become bactericidal, as it was the case for SEL-R1 and EMB-R5 (**Fig. 6**). Subinhibitory concentrations of selamectin, however, did not increase the activity of ethambutol in EMB-R11, which behaved as the parental strain. The role of EmbB in *M. smegmatis* susceptibility to selamectin was assessed by constructing an *embB* knockout strain in the mc^2^155 background and an overexpression strain. However, neither of them affected significantly the susceptibility to selamectin (**Table S7**).

**Fig. 6.**
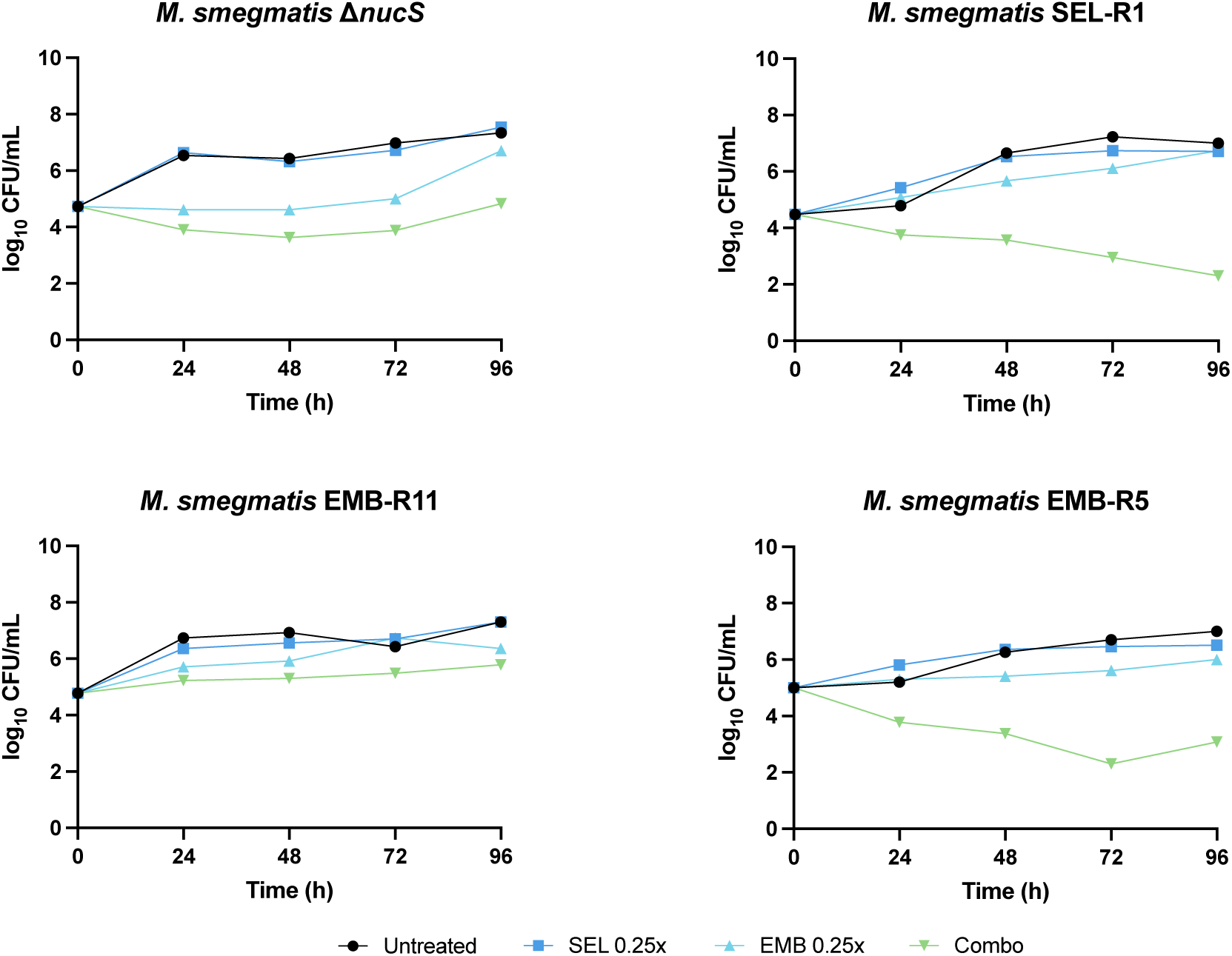
Time-kill kinetics of *M. smegmatis* mutants in the presence of combinations of selamectin and ethambutol at subinhibitory concentrations. Compounds were tested at 0.25-fold the MIC of the corresponding strain.

**Table 4.** Characterization of ethambutol-resistant mutants from the *M. smegmatis* SEL-R1 and Δ*nucS* strain backgrounds. Parental strains used for mutant isolation are marked with (*). Both parental strains were sequenced and showed no mutations when aligned to the reference *embB* sequence. SEL, selamectin.

| Background | Mutant name | EmbB mutation | MIC (μg/mL) | |
| --- | --- | --- | --- | --- |
|  |  |  | SEL | EMB |
| *M. smegmatis*<br>SEL-R1 | SEL-R1* | wild type | 16 | 2 |
|  | EMB-R2 | Y320H | 4 | 16 |
|  | EMB-R3 | G310R | 1 | 8 |
|  | EMB-R4 | Y320H | 8 | 16 |
|  | EMB-R5 | Q483R | 16 | 16 |
|  | EMB-R6 | Q483R | 16 | 16 |
|  | EMB-R7 | L388P | 2 | 16 |
|  | EMB-R8 | G310R | 1 | 4 |
|  | EMB-R9 | L388P | 2 | 16 |
| *M. smegmatis*<br>ΔnucS | ΔnucS* | wild type | 2 | 2 |
|  | EMB-R11, 13,<br>15, 16, 17, 18 | R307H | 2 | 16 |

## Discussion

In recent years, drug repurposing has emerged as a cost-effective alternative to the *de novo* drug discovery process. Following this strategy, a phenotypic screening identified the activity of the avermectins against *M. tuberculosis* and other mycobacteria [11–13]. Among avermectins, selamectin showed the best potency against mycobacteria. Selamectin is a veterinary avermectin used in cats and dogs, with improved pharmacokinetic properties and lower toxicity over other avermectins [30]. Understanding its mechanism of action can contribute to accelerate the progress through the drug discovery pipeline. In this study, we used mutant isolation approaches to study the mechanism of action of the avermectins against *M. smegmatis*, a non-pathogenic surrogate of *M. tuberculosis*.

Mutant isolation attempts with the wild-type *M. smegmatis,* and a transposon mutant pool allowed the isolation of colonies on high selamectin concentrations, but none of them displayed a stable resistant phenotype. This limitation was overcome using the mutator *M. smegmatis* Δ*nucS* strain, which allowed the isolation of intermediate selamectin resistant mutants (SEL-I with modest increases in their MIC) (**Fig.1**). WGS of SEL-I clones revealed the presence of nonsense mutations in two genes (*mmpL11* and *mps1*), both of which are involved in the biogenesis of the *M. smegmatis* envelope. GPLs are polar lipids exclusively found on the outer membrane of rapid-growing mycobacteria. Their presence or absence determines the overall hydrophobicity of the mycobacterial envelope, and therefore affects properties such as the ability to form biofilms. The lack of GPLs has been associated with aggregation and cording in fast-growing mycobacteria [31], which is consistent with the phenotype observed for the SEL-I mutants with nonsense mutations in the *mps1* gene. Analysis of the lipid profile of SEL-R mutants carrying the nonsense mutations in *mps1* confirmed that they were devoid of GPLs (**Fig. 4**). These contributions cannot be extrapolated to *M. tuberculosis*, which lacks GPLs. Deletion of *mmpL11* results in reduced membrane permeability, resistance to antimicrobial peptides, and alterations in biofilm formation. Its role seems to be conserved in *M. tuberculosis*, where it is required for full virulence in mice [25–27]. The introduction of both mutations in *M. smegmatis* Δ*nucS* reproduced the observed phenotype of the SEL-I strains (**Fig. 2**), confirming their contribution to the intermediate resistance phenotype. We hypothesize that the changes in the composition of the outer membrane could be affecting the permeability across the mycobacterial envelope, thus affecting selamectin susceptibility. The increased clumping tendency resulting from these mutations could promote reduction of selamectin susceptibility if bacteria growing within the clumps were exposed to lower selamectin concentrations (**Fig. S3**).

The SEL-I strains provided a genetic background to isolate high-level selamectin resistant mutants. WGS of SEL-R mutants provided a possible path for the development of the SEL-I phenotype: i.e., mutants derived from SEL-I2 acquired a missense mutation in *mmpL11*. It was not possible, however, to identify specific single mutations conferring high-level resistance. This was a limitation of the use of a mutator strain, where mutations conferring no adaptive advantages are more easily co-selected.

Selamectin resistance was reversed in the presence of subinhibitory concentrations of the cell wall inhibitors ethambutol, vancomycin, and cefradine (**Table 3**). Ethambutol targets the arabinosyltransferases EmbA and EmbB, involved in the synthesis of the arabinogalactan chains; and EmbC, involved in lipoarabinomannan biosynthesis [32,33]. Vancomycin and cefradine prevent peptidoglycan crosslinking by binding to the peptidoglycan pentapeptides, or by inactivating penicillin-binding proteins, respectively. The selamectin-ethambutol interaction was then studied in more depth using mutants with different susceptibilities to both drugs. Selection for ethambutol resistance using the SEL-R1 background yielded colonies that had reversed selamectin resistance. The EmbB Q483R mutation was the only mutation that did not restore selamectin susceptibility in SEL-R1-derived mutants. This mutation was the only one associated with clinical ethambutol resistance in *M. tuberculosis*, suggesting that mutations that restore selamectin susceptibility might have significant fitness costs on *M. smegmatis*. The fact that selamectin susceptibility can be restored by chemical or genetic inhibition of the mycobacterial envelope assembly suggests that mutations involved on the acquisition of high-level resistance are more likely impeding the access of selamectin to its target rather than affecting the interaction between selamectin and its target.

There are several cases in which selection of mutations on the target is not possible or does not lead to resistance that can be extrapolated to selamectin. First, selamectin could be acting on multiple targets. In this case, mutations in only one target would not confer resistance, owing that the other targets would still be susceptible. This would be consistent with our previous findings in which mutations in DprE1 (that altered the affinity of selamectin towards the enzyme) did not affect *M. smegmatis* susceptibility to selamectin [20]. Second, selamectin could be inhibiting multiple targets carrying out the same process. In this case, it is possible that the susceptible target dominates over the resistant one, as it occurs for aminoglycoside resistance in *E. coli* expressing different rRNA operon alleles [34], or bedaquiline susceptibility in strains expressing bedaquiline-susceptible and resistant *atpE* alleles simultaneously [35]. Third, the target might not be a protein and therefore its modification to become insensitive to selamectin requires the modification of a complete metabolic route, as it occurs with vancomycin, against which resistance can be achieved in *Staphylococcus aureus* by modifying the D-Ala-D-Ala moiety from peptidoglycan precursors but requires alternative enzymes that recognize the modified precursors [36].

Altogether, our results from mutant isolation assays and subsequent genetic and phenotypic characterization prove that the generation of high-level selamectin resistance is highly unlikely and requires multiple mutations to occur, which can be the result of selamectin acting on multiple targets. Being selamectin refractory to the generation of resistant mutants, this feature could be of great value in the development of novel regimens for TB treatment. Further studies are needed to understand the mode of action of selamectin against the pathogenic *M. tuberculosis* and how to best use it in combinatorial regimens.

## Materials and methods

### Bacterial strains, compounds, and culture conditions

*M. smegmatis* strains were routinely grown at 37 °C in Middlebrook 7H9 broth (Difco) supplemented with 10% Middlebrook Albumin-Dextrose-Catalase (ADC) (Difco) and 0.05% (v/v) Tyloxapol or on Middlebrook 7H10 agar (Difco) supplemented with 10% (v/v) Middlebrook Oleic Acid-Albumin-Dextrose-Catalase (OADC) (Difco). For strains carrying selectable markers, antibiotics were added at the following concentrations: kanamycin (Km, Sigma), 20 µg/mL; hygromycin B (Hyg, Invivogen); 20 µg/mL, streptomycin (Sm, Sigma), 20 µg/mL. Selamectin was purchased from the European Pharmacopoeia.

### Drug susceptibility assays

Minimal inhibitory concentrations (MIC) were determined in 7H9 broth supplemented with 0.2% glycerol and 10% ADC, using two-fold serial dilutions of the compounds in duplicates in 96-well flat-bottomed polystyrene plates. Plates were inoculated with 10^5^ CFU/mL of *M. smegmatis* and incubated in the presence of the compounds for 3 days before the addition of 30 µL of a solution of the bacterial growth reporter MTT [3-(4,5-dimethylthiazol-2-yl)-2,5-diphenyltetrazolium bromide] (2.5 mg/mL) and Tween 80 (10%, v/v). Following a 3-hour incubation, the optical density at 580 nm (OD_580_) was measured, and the MIC was defined as the lowest concentration of compound that inhibited at least 90% of MTT conversion to formazan.

*M. smegmatis* MICs were also determined on 7H10 agar with 10% OADC, testing two-fold serial dilutions of the compounds in duplicates in 24-well polystyrene plates. Compounds were added to tempered agar and mixed thoroughly in 15 mL centrifuge tubes before transferring 2 mL to each well. Three different inoculums: 10^3^, 10^5^, and 10^7^ total CFU, were seeded onto each well by transferring 10 µL of bacterial suspension and incubated for 3 days. The MIC was defined as the minimal concentration of compound which inhibited colony formation.

### Checkerboard assays

Checkerboard assays were performed as previously described [37]. Compound stocks were prepared at 4x the highest test concentration for each compound. Then, the two compounds were diluted by making two-fold dilutions across all rows and columns, respectively. Plates were inoculated with 10^5^ CFU/mL, and the readout was performed as described for MIC determinations. The fractional inhibitory concentrations (FIC) were defined as the ratio between the MIC of either compound in the combination and the MIC of the compound alone. Then, FIC indices (FICI) were obtained by adding each drug FIC, and the lowest FICI value was used to determine whether an interaction was synergistic (FICI ≤ 0.5), there was no interaction (0.5 < FICI < 4) or was antagonistic (FICI ≥ 4) [38].

### Time-kill kinetic assays

25 cm^2^ culture flasks containing 10 mL of 7H9 with 0.2% glycerol (v/v) and 10% ADC were inoculated with *M. smegmatis* to a final density of 10^5^ CFU/mL from liquid cultures in late exponential phase. Aliquots of the culture were taken at 0, 6, 24, 48, 72, and 96 hours following the addition of the compounds; at each timepoint, 10-fold serial dilutions were made in PBS with 0.1% tyloxapol and 100 µL were seeded onto Luria-Bertani (LB) agar plates. Colony-forming units (CFUs) were determined after 3 days of incubation at 37 °C.

### Selection of selamectin-resistant mutants

Mutant isolation was first attempted by plating 10^5^ to 10^8^ total CFUs of *M. smegmatis* mc^2^155 or a transposon library made in *M. smegmatis* HS42 using the TnSPAZ transposon [21] onto 90 mm Petri dishes containing 1, 2, 4, 10, or 20-fold the MIC of the selamectin against the corresponding inoculum adjusted per surface unit. Because the surface of a 90 mm Petri dish is roughly 100 times the surface of a 10 µL droplet, the MIC value chosen as reference for each inoculum was the one corresponding to a 100-fold lower inoculum in a 24-well plate.

Mutant isolation attempts with the *M. smegmatis* Δ*nucS* strain [23] were performed under conditions that restricted *M. smegmatis* mc^2^155 growth (4 to 20-fold the MIC and inoculums below 10^7^ CFUs). Two of these colonies (SEL-I1 and SEL-I2) were subsequently used for the isolation of high-level resistance on plates containing 10 or 20-fold the MIC of selamectin.

In all cases, plates were incubated at 37 °C for up to 7 days. For their phenotypic validation, the colonies isolated were transferred to drug-free medium and their susceptibility to selamectin was determined using three different assays. First, the MIC of selamectin against each strain was determined as above described but using 1.4-fold dilution steps instead of conventional two-fold dilution steps. Second, susceptibility to selamectin was tested on agar plates by seeding 5 µL spots of four 10-fold serial dilutions (starting from 10^7^ bacteria/mL, i.e., 5·10^4^ total bacteria) onto plates containing 0, 1, 4, 16, or 40 µg/mL of selamectin. In a final validation step, resistance to selamectin was confirmed by time-kill kinetic assays. Those clones with decreased selamectin susceptibility were selected for WGS.

### Selection of ethambutol-resistant mutants

Ethambutol-resistant mutants were selected by plating approximately 10^7^ CFUs of *M. smegmatis* onto 90 mm 7H10-OADC plates containing 4-fold the MIC of ethambutol on solid media. The colonies isolated were then transferred to drug-free medium and their resistant phenotype confirmed by MIC determination in liquid medium.

### Whole genome sequencing and identification of polymorphisms

Genomic DNA from the resistant *M. smegmatis* mutants was extracted using the CTAB (Cetyl Trimethyl Ammonium Bromide) method [39]. Briefly, bacteria were resuspended in 400 µL TE (100 mM Tris/HCl, 10 mM EDTA, pH = 8.0) and heated at 85 °C for 10 min. Samples were then chilled and treated with 50 µL of lysozyme (10 mg/mL) at 37 °C for at least 1 hour. Hereafter, 72.5 µL of 10% sodium dodecyl sulfate and 2.5 µL of proteinase K (20 mg/mL) were added and the resulting mixture was incubated at 65 °C for 10 min. Subsequently, 100 µL of 5 M NaCl and 100 µL of prewarmed CTAB/NaCl (10% CTAB in 0.7 M NaCl) were added and incubated at 65 °C for a further 10 min. Genomic DNA was extracted by adding 750 µL of chloroform:isoamyl alcohol (24:1). Samples were centrifugated (5 min, 12,000 RCF) and the upper phase was transferred to a fresh tube containing 450 µL isopropanol. Nucleic acids were precipitated overnight at -20 °C and were collected by centrifugation (10 min, 12,000 RCF). The pellets were dissolved in 50 µL nuclease-free water and quantified using an ND-1000 spectrophotometer.

Genomic DNA was sequenced at the FISABIO Sequencing and Bioinformatics Service (Valencia, Spain). DNA libraries were prepared using the Nextera XT DNA Library Prep kit (Illumina) following manufacturer’s instructions and sequenced on the Illumina MiSeq platform. The Illumina paired-end reads were filtered with fastp [40] to remove low-quality bases at the 3’ end. The filtered reads were subsequently mapped to the *M. smegmatis* chromosome (GenBank accession number CP001663.1) with BWA [41] and potential duplicates were removed with Picard tools (http://broadinstitute.github.io/picard). Single nucleotide polymorphisms (SNPs) were called with VarScan [42] if at least 20 reads supported the genomic position, the SNP was found at a frequency of 0.9 or higher and was not found near an indel region (10 bp). Indels were called with Genome Analysis ToolKit (GATK) [43]. SNPs and indels were annotated by using SnpEFF [44], and those common to the parental strain were later removed.

### Genetic manipulation of *M. smegmatis*

*M. smegmatis* single-point mutants were constructed using single-stranded DNA recombineering [45]. Mutagenic oligonucleotides were designed so that they would introduce additional silent mutations to generate a new restriction site (**Table S8**).

Recombineering-induced *M. smegmatis* were prepared from cultures grown to mid-exponential phase (OD_600_ = 0.5). At that point, acetamide was added to a final concentration of 0.2% (w/v), and the culture was incubated for an additional 4 hours to allow induction of the recombineering system. Bacteria were then transferred to ice for 30 min, harvested (5 min, 4,000 RCF), washed four times with ice-cold 10% glycerol, and resuspended in 1/100 of the initial culture volume.

Recombineering-induced *M. smegmatis* Δ*nucS* pJV53H cells were electroporated in 0.2 cm cuvettes (Bio-Rad) and applied a 2500 V, 25 µF, 1000 Ω pulse in a GenePulser Xcell electroporator (Bio-Rad) with a mixture of 500 ng of the oligonucleotide carrying the desired mutations and 100 ng of an oligo targeting *rpsL* that confers Sm resistance. Bacteria were then recovered in 1 mL of fresh medium, incubated overnight at 37 °C, and selected on LB agar plates containing 20 µg/mL Sm. Colonies were screened for the presence of the desired point mutation by colony PCR after digestion of the PCR products with either *Xba*I or *Bsp*TI (Thermo Fisher) for one hour. Positive colonies, containing a mixture of bacteria carrying wild-type and mutant alleles, were streaked onto LB agar plates to isolate isogenic colonies, which were then rescreened using the same methodology.

Multiple mutations were introduced using 500 ng of pBP10 plasmid [46] for co-selection instead of the oligonucleotide conferring Sm resistance. Km-resistant colonies were screened as above described for single point mutations. Finally, once a colony with the desired mutation was obtained, pBP10 was cured by doing a passage in medium without Km and seeding bacteria onto LB agar plates. Curation was verified using PCR with oligos targeting the Km resistance cassette.

### Analysis of *M. smegmatis* lipids and mycolic acids by thin-layer chromatography (TLC)

Cells were grown by shaking at 37 °C in 7H9 broth supplemented with 10 % Albumin-Dextrose-Catalase in the presence or absence of 0.05 % Tween80. After cultivation, cells were harvested by centrifugation (2200 x RCF, 20 min). Lipids were extracted from cell pellets by extraction with 6 mL chloroform/methanol (1:2) at 56 °C for 2 h, followed by two extractions in 6 mL chloroform/methanol (2:1) under the same conditions. Extracts were pooled into a glass tube, dried under a stream of nitrogen, and subjected to biphasic wash in chloroform/methanol/water (4:2:1), as described by Folch [47]. After washing, organic phases from each sample were dried and dissolved in 600 µL chloroform/methanol (2:1) per 0.5 g of wet cell weight. Then, 10 µL of each sample were loaded on thin-layer chromatography (TLC) silica gel 60 F_254_ plates (Merck) and lipids were separated in following eluents - solvent I (chloroform/methanol/water 20:4:0.5); solvent II (chloroform/methanol/ammonium hydroxide/water 65:25:0.5:4) and solvent III (petroleum ether/ethyl acetate 95:5; 3 runs). Lipids were visualized by soaking the plates in 10 % CuSO_4_ in 8 % phosphoric acid solution and charring with heat gun.

For isolation of glycopeptidolipids (GPLs), 50 µL from obtained lipid extracts were dried and subjected to alkaline methanolysis (200 µL 0.2 M NaOH in methanol for 2 h at 37 °C) to cleave ester-linked fatty acids [48]. After pH neutralization with glacial acetic acid, samples were dried under nitrogen flow and alkaline-stable GPLs were partitioned by a biphasic system composed of chloroform/methanol/water (4:2:1) [47]. Organic phase was washed twice with chloroform/methanol/water (3:48:47), dried under the stream of nitrogen and resuspended in 50 µL chloroform/methanol (2:1). Then, 10 µL from purified GPLs were applied on silica gel 60 F_254_ plates, separated in solvent IV (chloroform/methanol 9:1) and visualized by soaking the plates in solution of orcinol in sulfuric acid followed by charring.

Methyl esters of mycolic acids (MAME) and fatty acids (FAME) were prepared from whole cells as previously described [49]. Briefly, 2 ml of 15 % tetrabutylammonium hydroxide solution were added to pelleted cells and samples were saponified overnight at 100 °C. After cooling down, samples were derivatized to corresponding methyl esters by 4 h incubation in the mixture of 3 mL dichloromethane, 300 µL iodomethane and 1 mL water at room temperature on a rotary shaker. Methylated samples were washed with water three times and resulting organic phase was dried under nitrogen flow. MAME and FAME were extracted to 3 mL diethyl ether and dried. Samples were dissolved in chloroform/methanol (2:1) and loaded on TLC plates as described for lipid extracts. Different forms of methyl esters were separated in solvent V (*n*-hexane/ethyl acetate 95:5; 3 runs) and visualized as previously stated for lipids.

## Acknowledgements

We thank Katarína Mikušová from the Comenius University in Bratislava for helpful discussions during the performance of this work; and Ana Picó and Begoña Gracia from Universidad de Zaragoza for technical support. Porin-deficient *M. smegmatis* strains were a gift from Michael Niederweis.

## Authors’ Contributions

CRediT (Contributor Roles Taxonomy) has been applied for author contribution. Conceptualization: J.M.E.-A., S.R.-G.; Investigation: J.M.E.-A., H.G., J.K.; Formal Analysis: J.M.E.-A., H.G., A.C.-O.; Resources: A.L., J.B.; Supervision: A.L, J.K., I.C., J.A.A., S.R.-G., Writing – original draft: J.M.E.-A., H.G., J.K. Writing – review and editing: J.M.E.-A., H.G., J.K, J.A.A., S.R.-G. All authors have reviewed and agreed to the final version of the manuscript.

---

**Fig. S1.**
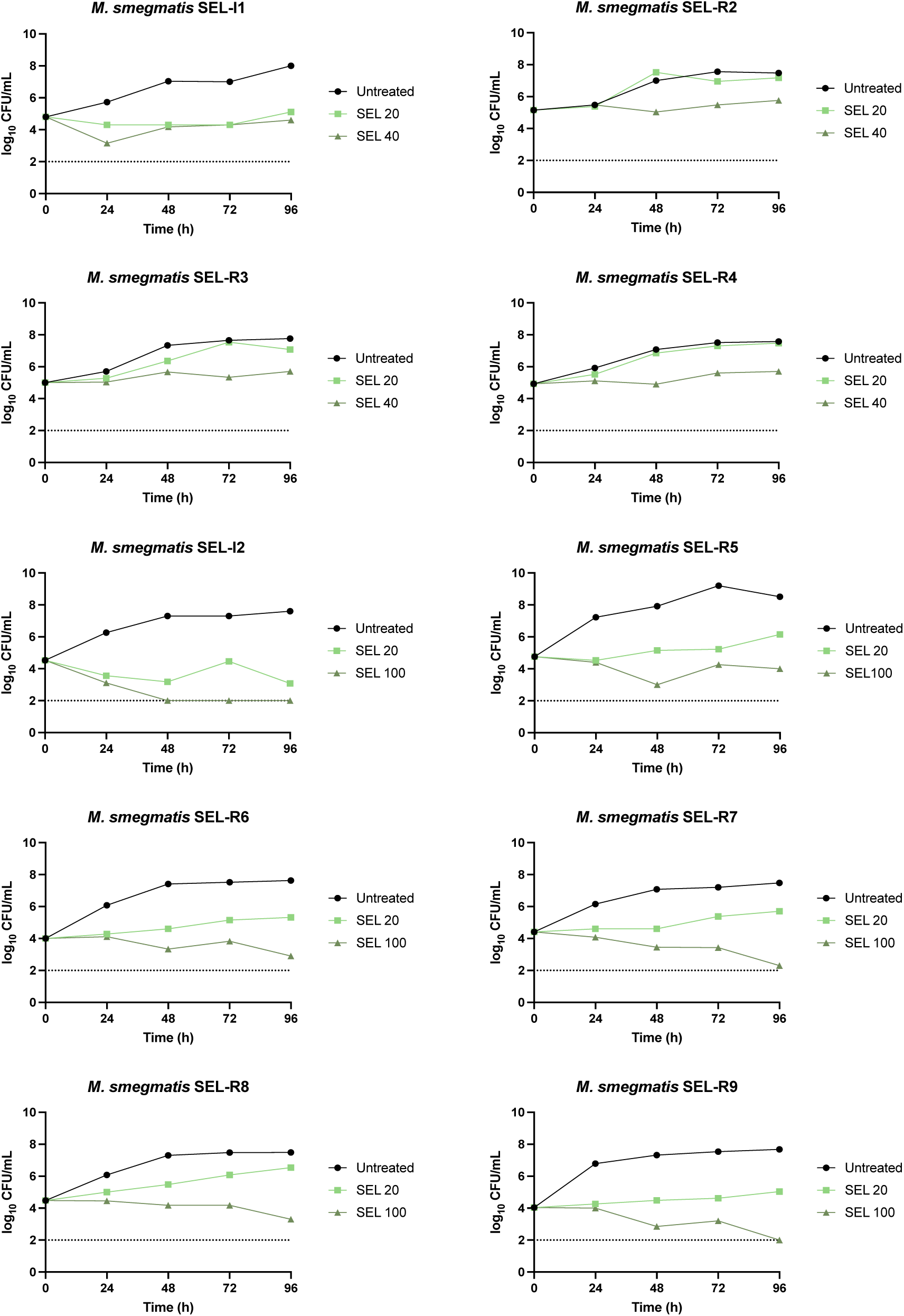
Time-kill kinetics of selamectin SEL-R clones, and their parental strains. SEL-R(2-4) derived from SEL-I1, SEL-R(5-9) derived from SEL-I2. Selamectin (SEL) concentrations are given in µg/mL. Limit of detection: 100 CFU/mL (dotted line).

**Fig. S2.**
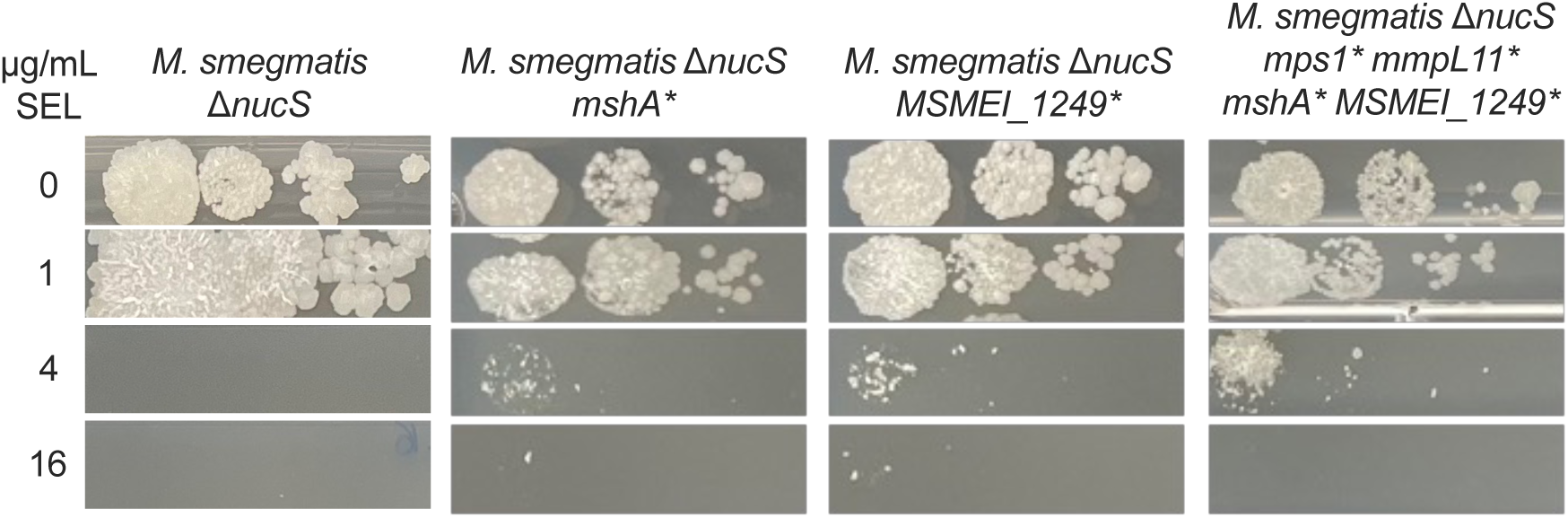
Phenotypic validation of *M. smegmatis* engineered point mutants at *mshA* and *MSMEI_1249* by culture drop dilution in plates containing different selamectin concentrations (expressed in µg/mL). Ten-fold serial dilutions of the bacterial inoculum were made from left to right. *: truncated protein, due to a stop codon in the ORF. SEL, selamectin.

**Fig. S3.**
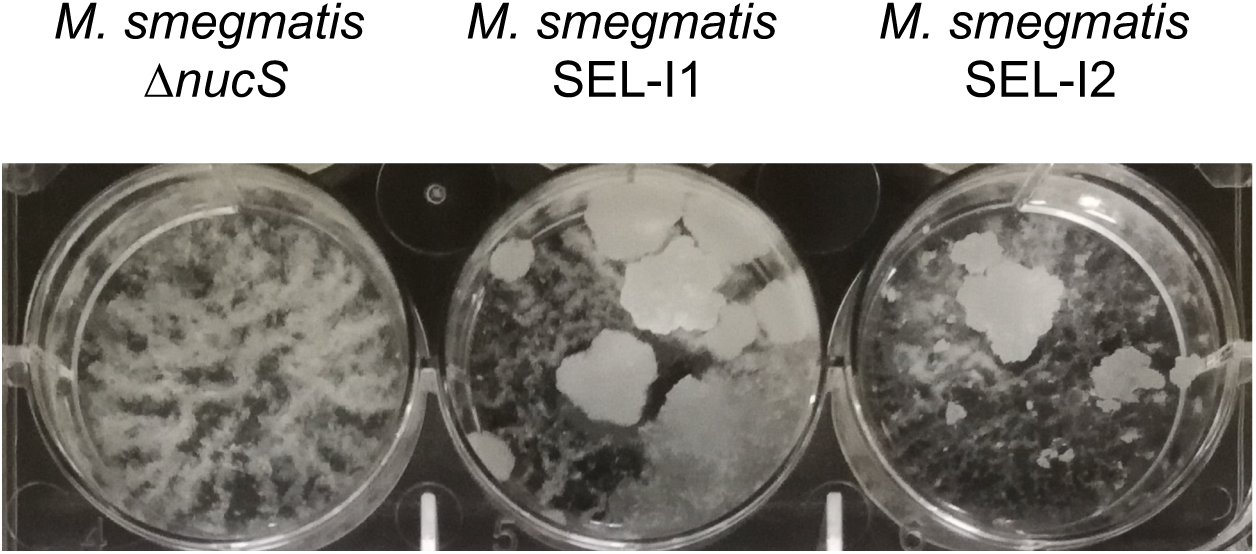
*M. smegmatis* SEL-I mutants have an increased clumping tendency. Strains were grown in 7H9-0.2% glycerol-ADC.

**Table S1.** Isolation of low-level selamectin resistant mutants. The mutator *M. smegmatis* Δ*nucS* strain was exposed to selamectin at different cell densities. Mutant frequencies were calculated as the ratio between the number of colonies isolated and the total CFU plated. SEL, selamectin; *The inoculum was split in 10 plates

| Strain | Inoculum | Compound concentration (x MIC) | Number of colonies isolated | Mutation frequency |
| --- | --- | --- | --- | --- |
| *M. smegmatis* ΔnucS | 7.4·10<sup>4</sup> CFU | SEL 16 µg/mL (4x) | 0 | - |
|  |  | SEL 40 µg/mL (10x) | 0 | - |
|  |  | SEL 80 µg/mL (20x) | 0 | - |
|  | 7.4·10<sup>5</sup> CFU | SEL 16 µg/mL (4x) | 1 | 1.4·10<sup>-6</sup> |
|  |  | SEL 40 µg/mL (10x) | 0 | - |
|  |  | SEL 80 µg/mL (20x) | 0 | - |
|  |  | INH 160 µg/mL (10x) | 386 | 5.2·10<sup>-4</sup> |
|  | 7.4·10<sup>6</sup> CFU | SEL 40 µg/mL (10x) | 10 | 1.4·10<sup>-6</sup> |
|  |  | SEL 80 µg/mL (20x) | 2 | 2.7·10<sup>-7</sup> |
|  | 1.5·10<sup>8</sup> CFU* | SEL 40 µg/mL (10x) | 25 | 1.7·10<sup>-7</sup> |
| *M. smegmatis* mc<sup>2</sup>155 | 10<sup>6</sup>-10<sup>8</sup> CFU | SEL 16-80 µg/mL (4x-20x) | 0 | - |
|  | 1·10<sup>6</sup> CFU | INH 160 µg/mL (10x) | 18 | 1.8·10<sup>-5</sup> |

**Table S2.** Non-synonymous mutations identified in intermediate selamectin-resistant mutants. Amino acid mutations for each locus are denoted in parentheses. Frameshift mutations are labeled as *fs*; insertions, as *ins*; and deletions, as *del*.

| Locus | Predicted function | Strain |
| --- | --- | --- |
| *mmpL11* (Q452-STOP) | Membrane transporter MmpL11 | I1 |
| *mps1* (Q3256-STOP) | Non-ribosomal peptide synthase | I1, I2, I3, I4, I5 |
| *mps2* (G1816E) | Non-ribosomal peptide synthase | I9 |
| *MSMEI_1397* (E78G) | Uncharacterized protein | I1 |
| *MSMEI_1646* (L192fs) | NPL/P60-family secreted protein | I4 |
| *gpsI* (T759A) | Polyribonucleotide nucleotidyl transferase | I1 |
| *MSMEI_2709* (Q395R) | Ribonuclease D | I1 |
| *MSMEI_2715* (R86Q) | Peptide-methionine (R)-S-oxide reductase | I1, I2, I3, I4, I5 |
| *MSMEI_3478* (G252D) | Short-chain dehydrogenase | I9 |
| *MSMEI_4034* (P289fs) | Transposase, IS4 family protein | I4, I6, I8, I9, I10 |
| *MSMEI_4108* (E1201G) | Putative nitrate reductase | I1 |
| *MSMEI_4508* (R74fs) | 50S ribosomal protein L21 | I8 |
| *MSMEI_5940* (L274insLL;<br>E277_Q278del) | Uncharacterized protein | I6 |
| *MSMEI_6237* (G78S) | Erp protein | I9 |
| *MSMEI_6491* (E307G) | Sulfate/thiosulfate importer (ABC transporter) | I1 |
| *MSMEI_6509* (L147P) | Purine catabolism PurC regulatory protein | I1, I2, I4 |

**Table S3.** Non-synonymous mutations identified in SEL-R mutants isolated from the SEL-I1 background (*mmpL11* and *mps1* nonsense mutations). MSH: mycothiol; *ins*: insertion.

| Locus | Predicted function |
| --- | --- |
| **MSMEI_0910 (*mshA*) (135-STOP)** | D-inositol-3-phosphate transferase. Involved in MSH biosynthesis |
| *MSMEI_1146* (S229P) | Putative allantoin permease |
| *MSMEI_1197* (E295G) | Nitrate ABC transporter |
| **MSMEI_1249 (35-STOP)** | Putative polyketide cyclase/dehydrase |
| *MSMEI_1333* (S155G) | Sugar ABC transporter |
| *MSMEI_1942* (G340R) | D-malate dehydrogenase |
| *MSMEI_3032* (L377P) | Extracellular substrate-binding protein |
| *MSMEI_3447* (E13G) | *gntR* family transcription factor |
| *MSMEI_3477* (V221A) | Putative dienelactone hydrolase |
| *MSMEI_4923* (W260R) | MFS transporter |
| *MSMEI_4999* (L378P) | Uncharacterized protein |
| *MSMEI_5975* (A75V) | Predicted aldehyde dehydrogenase |
| *MSMEI_6346* (Q111R) | Orotate phosphoribosyltransferase PyrE |
| *MSMEI_4555* (*clpX*) (-7insG) | ClpX ATPase subunit. |

**Table S4.**
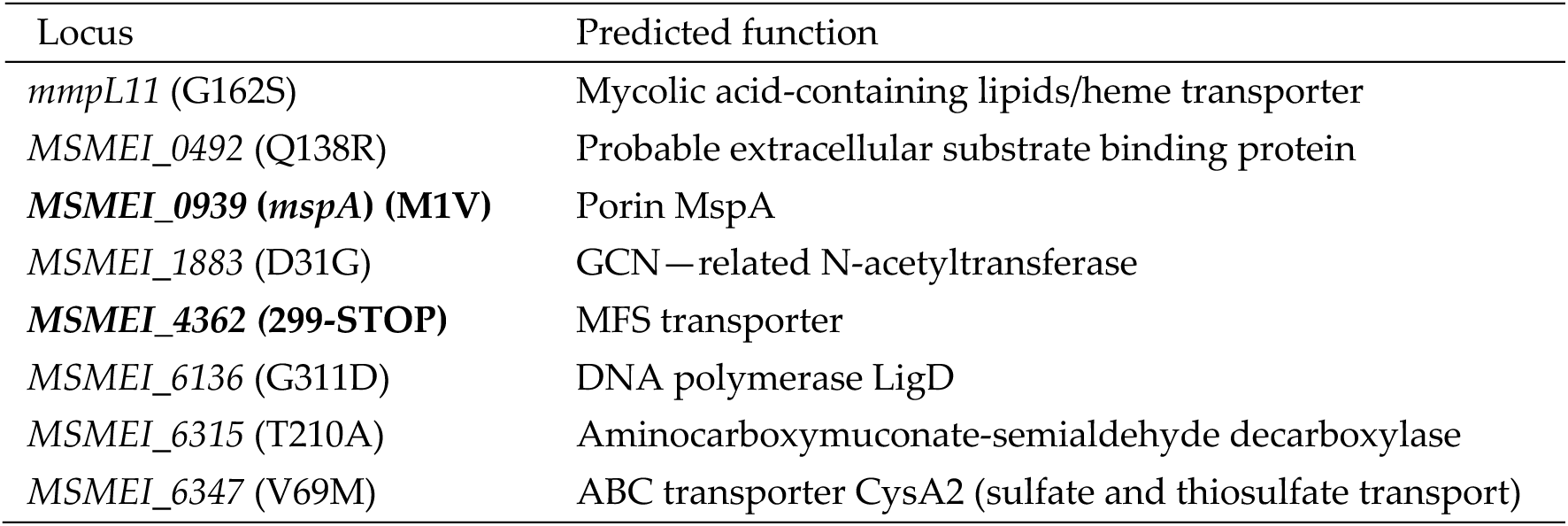
Common non-synonymous mutations identified in SEL-R mutants isolated from SEL-I2 background (*mps1* nonsense mutations). Mutations common to four of the five mutants are displayed. The SEL-R7 mutant only had the clpX (-7insG) mutation.

**Table S5.**
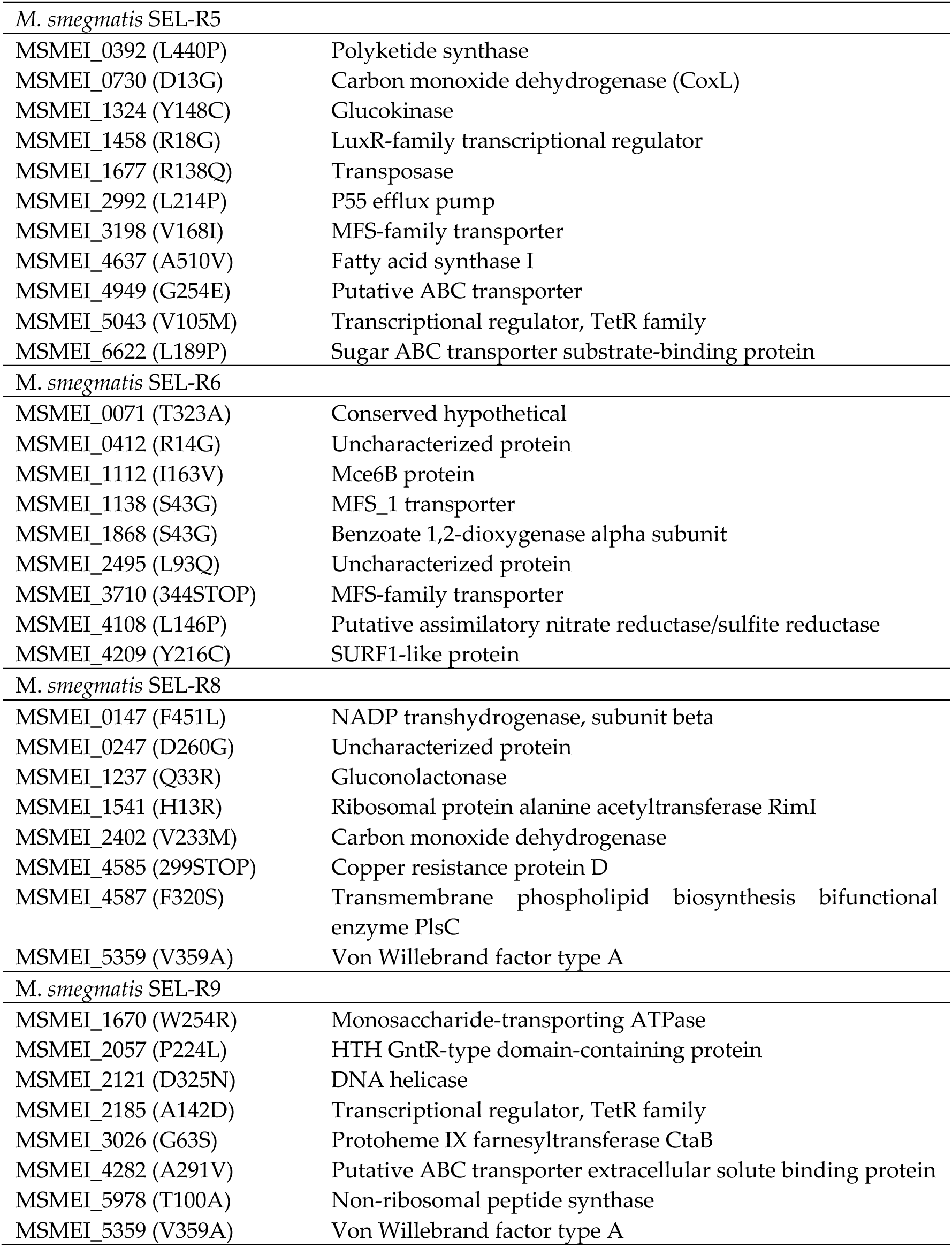
Mutant-specific non-synonymous mutations identified in SEL-R mutants isolated from SEL-I2 background (*mps1* nonsense mutations).

|  |  |
| --- | --- |
| *M. smegmatis* SEL-R5 |  |
| MSMEI_0392 (L440P) | Polyketide synthase |
| MSMEI_0730 (D13G) | Carbon monoxide dehydrogenase (CoxL) |
| MSMEI_1324 (Y148C) | Glucokinase |
| MSMEI_1458 (R18G) | LuxR-family transcriptional regulator |
| MSMEI_1677 (R138Q) | Transposase |
| MSMEI_2992 (L214P) | P55 efflux pump |
| MSMEI_3198 (V168I) | MFS-family transporter |
| MSMEI_4637 (A510V) | Fatty acid synthase I |
| MSMEI_4949 (G254E) | Putative ABC transporter |
| MSMEI_5043 (V105M) | Transcriptional regulator, TetR family |
| MSMEI_6622 (L189P) | Sugar ABC transporter substrate-binding protein |
| *M. smegmatis* SEL-R6 |  |
| MSMEI_0071 (T323A) | Conserved hypothetical |
| MSMEI_0412 (R14G) | Uncharacterized protein |
| MSMEI_1112 (I163V) | Mce6B protein |
| MSMEI_1138 (S43G) | MFS_1 transporter |
| MSMEI_1868 (S43G) | Benzoate 1,2-dioxygenase alpha subunit |
| MSMEI_2495 (L93Q) | Uncharacterized protein |
| MSMEI_3710 (344STOP) | MFS-family transporter |
| MSMEI_4108 (L146P) | Putative assimilatory nitrate reductase/sulfite reductase |
| MSMEI_4209 (Y216C) | SURF1-like protein |
| *M. smegmatis* SEL-R8 |  |
| MSMEI_0147 (F451L) | NADP transhydrogenase, subunit beta |
| MSMEI_0247 (D260G) | Uncharacterized protein |
| MSMEI_1237 (Q33R) | Gluconolactonase |
| MSMEI_1541 (H13R) | Ribosomal protein alanine acetyltransferase RimI |
| MSMEI_2402 (V233M) | Carbon monoxide dehydrogenase |
| MSMEI_4585 (299STOP) | Copper resistance protein D |
| MSMEI_4587 (F320S) | Transmembrane phospholipid biosynthesis bifunctional enzyme PlsC |
| MSMEI_5359 (V359A) | Von Willebrand factor type A |
| *M. smegmatis* SEL-R9 |  |
| MSMEI_1670 (W254R) | Monosaccharide-transporting ATPase |
| MSMEI_2057 (P224L) | HTH GntR-type domain-containing protein |
| MSMEI_2121 (D325N) | DNA helicase |
| MSMEI_2185 (A142D) | Transcriptional regulator, TetR family |
| MSMEI_3026 (G63S) | Protoheme IX farnesyltransferase CtaB |
| MSMEI_4282 (A291V) | Putative ABC transporter extracellular solute binding protein |
| MSMEI_5978 (T100A) | Non-ribosomal peptide synthase |
| MSMEI_5359 (V359A) | Von Willebrand factor type A |

**Table S6.** MIC of selamectin for *M. smegmatis* porin deficient mutants. *M. smegmatis* SMR5 is the parental strain that was used to engineer the mutants.

| Strain | MIC (µg/mL) |
| --- | --- |
| *M. smegmatis* SMR5 (parental strain) | 4 |
| *M. smegmatis* MN01 ($\Delta mspA$) | 4 |
| *M. smegmatis* ML10 ($\Delta mspA \Delta mspC$) | 4 |

**Table S7.** MICs of selamectin, ethambutol, and rifampicin against *embB* knockout and overexpression mutants. MIC values are given in µg/mL.

| Strain | Selamectin | Ethambutol | Rifampicin |
| --- | --- | --- | --- |
| *M. smegmatis* mc<sup>2</sup>155 | 2,83 | 1 | 64 |
| *M. smegmatis* mc<sup>2</sup>155 $\Delta embB$ | 2 | 1 | 0,25 |
| *M. smegmatis* mc<sup>2</sup>155 pMV261-*embB* | 2 | 1 | 64 |

**Table S8.**
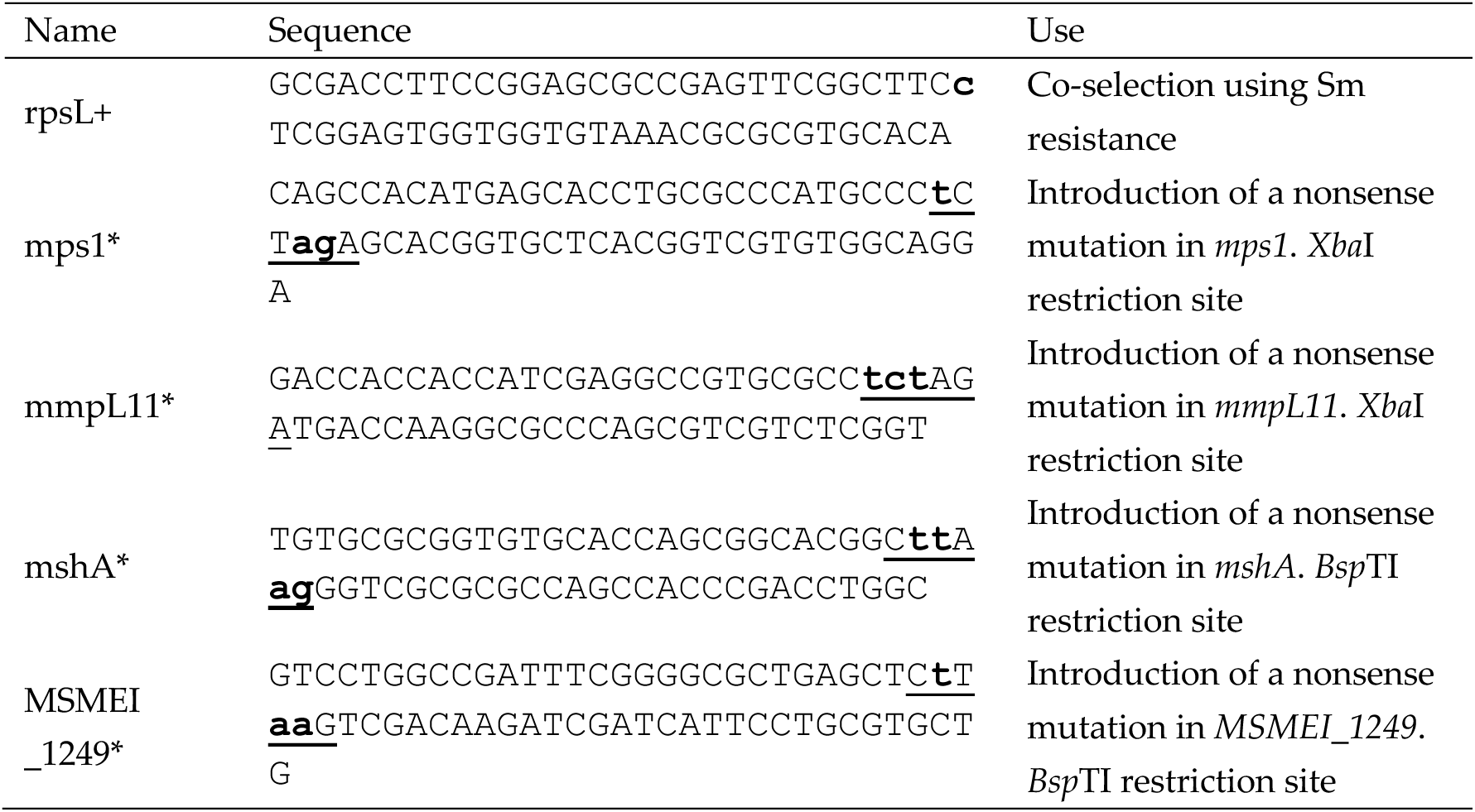
Mutagenic oligonucleotides used for ssDNA recombineering. Bold lowercase characters show point mutations introduced by the oligonucleotides; restriction sites generated by the point mutations are underlined.

